# Usp14 is required for spermatogenesis and ubiquitin stress responses in *Drosophila melanogaster*

**DOI:** 10.1101/724153

**Authors:** Levente Kovács, Ágota Nagy, Margit Pál, Peter Deák

## Abstract

Deubiquitinating (DUB) enzymes free covalently linked ubiquitins from ubiquitin-ubiquitin and ubiquitin-protein conjugates, and thereby maintain the equilibrium between free and conjugated ubiquitins and regulate ubiquitin-mediated cellular processes. The present genetic analyses of mutant phenotypes demonstrate that loss of Usp14 function results in male sterility, with defects in spermatid individualization and reduced testicular free monoubiquitin levels. These phenotypes were rescued by germline specific overexpression of wild type Usp14. Synergistic genetic interactions with *Ubi-p63E* and cycloheximide sensitivity suggest that ubiquitin shortage is a primary cause of male sterility. In addition, *Usp14* is predominantly expressed in testes in *Drosophila*, and differential expression patterns may be causative of testis-specific loss of function *Usp14* phenotypes. Collectively, these results suggest a major role of Usp14 in maintaining normal steady state free monoubiquitin levels during the later stages of *Drosophila* spermatogenesis.

## INTRODUCTION

Deubiquitinating enzymes (DUBs) are ubiquitin-specific proteases that hydrolyze isopeptide bonds between ubiquitin–ubiquitin and ubiquitin–protein conjugates, and thereby act as key regulators of ubiquitin-dependent processes (Amerik and Hochstrasser, 2004). Opposing ubiquitination and deubiquitination activities maintain a dynamic intracellular equilibrium between free and conjugated ubiquitin forms, in which monomeric ubiquitins are indispensable for normal cell physiology and development. About 100 DUB genes have been identified in humans, and these are classified into five or six conserved families of DUBs according to their catalytic domains (Nijman et al., 2005; Suresh et al., 2016). Basic DUB activities include processing of ubiquitin precursors and ubiquitin recycling following deubiquitination and/or polyubiquitin chain editing (Kim et al., 2003; Reyes-Turcu et al., 2009).

The evolutionarily conserved proteasome associated DUBs Rpn11, Uch-L5, and Usp14 have prominent roles in protein recycling, and release ubiquitin from proteasome bound substrate proteins prior to translocation and proteasomal degradation (Guterman and Glickman, 2004). Binding to the proteasome was previously associated with disassembling activities of Ubp6, the yeast orthologue of Usp14, and deletion of this protein led to reduced monoubiquitin levels (Legget et al., 2002). Several pleiotropic effects of Usp14/Ubp6 loss have been identified, including synaptic transmission defects, ataxia, and premature death in mice (Wilson et al., 2002). Dual roles of Ubp6 in regulating ubiquitin levels and proteasome function were described in yeast (Hanna et al., 2007), and loss of this protein reduced the yeast prion [PSI^+^] expression and heightened sensitivity to a broad range of toxic drugs (Chernova et al., 2003). *Usp14* inactivation delayed cell proliferation of mouse embryonic fibroblasts and in the developing Drosophila eye (Lee et al., 2018). Moreover, gross perturbations of ubiquitin equilibria and reduced free monoubiquitin pools are common to all of these Usp14/Ubp6 related abnormalities.

In our studies of Usp5 in Drosophila, deficiencies of free monoubiquitins triggered ubiquitin stress responses, which were correlated with increased Usp14 deubiquitinase expression (Kovács et al., 2015). To investigate biological functions, loss of function *Usp14* mutants were isolated and subjected to phenotypic characterization. Our male *Usp14* null mutants were sterile and had sperm individualization defects and low free monoubiquitin levels in the testes. Reduced ubiquitin levels in these mutants was confirmed by high sensitivity to the translational inhibitor cycloheximide and an additive phenotype of *Usp14*–*Ubi-p63E* double mutants. These phenotypes are consistent with predominant Usp14 expression in testes.

## RESULTS AND DISCUSSION

### *Usp14* mutant males are sterile

The *Drosophila* orthologue of Usp14 is encoded by the *CG5384* gene and has 49% overall similarity with yeast Ubp6 and 71% similarity with both mouse and human USP14 ubiquitin proteases. Catalytic ubiquitin-specific protease and proteasome binding ubiquitin-like (UBL) domains of these proteins are highly conserved (Tsou et al., 2012; Kovács et al., 2015). Our previous study of *Drosophila* orthologue of Usp5 showed increased expression of *Usp14* in *DmUsp5* mutants, suggesting roles of Usp14 in ubiquitin stress responses to ubiquitin shortages (Kovács et al., 2015), as described in yeast (Hanna et al., 2007).

To investigate the physiological consequences of Usp14 depletion, we analyzed phenotypes following PiggyBac insertion of a *Usp14* allele (*Usp14*^*f*^). The PiggyBac element was inserted between the first intron and second exon (Fig. 1A) and resulted in substantial reductions in *Usp14* mRNA expression levels, indicating that *Usp14*^*f*^ is a relatively strong hypomorph allele (Fig. 1B). Although *Usp14*^*f*^ homozygotes were viable, all eclosed males were sterile (Fig. 1C, column 3) and despite normal mating behaviors, their wild type female mating partners laid mostly unfertilized eggs. In phase contrast and fluorescent microscopy analyses of *Usp14*^*f*^ homozygote testes, all stages through spermatid elongation were indistinguishable from those in wild type flies. Although sperm bundles were formed in *Usp14*^*f*^ testes, seminal vesicles were almost empty and contained few individualized sperms (Fig. 1D). In addition, most sperm bundles were accumulated before the entrance of seminal vesicles, suggesting defects in sperm individualization. Male sterility was also observed when *Usp14*^*f*^ was situated *in trans* of a deficiency uncovering the *Usp14* locus (Fig.1C, column 5). Yet, male fertility was completely restored in revertants of *Usp14*^*f*^ (Fig. 1C, column 2). Viability and fertility of females were not affected by *Usp14* mutations (data not shown).

**Fig. 1.**
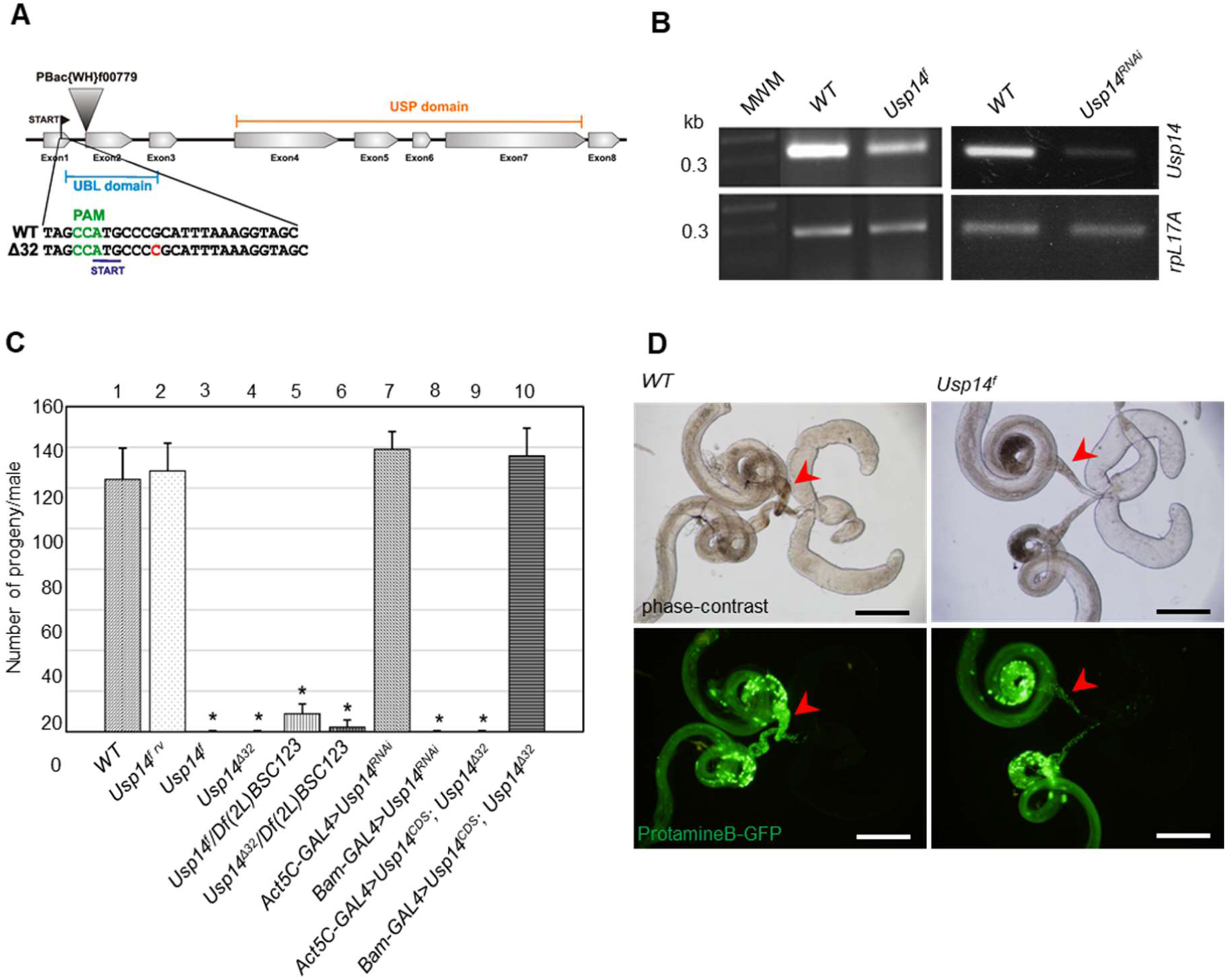
Male sterile phenotype of *Usp14* mutants; (A) diagram of the Usp14 gene; arrowhead shows the position of the PiggyBac insertion (denoted as *Usp14*^*f*^ in text). Insertion of single cysteine in *Usp14*^*Δ32*^ mutants is highlighted in red. (B) *Usp14* expression levels in mutant and knockdown flies were determined using semi quantitative reverse transcriptase-polymerase chain reaction (RT-PCR) analyses of wild type, mutant, and RNAi transfected flies. *Usp14*^*RNAi*^ was controlled by the ubiquitous *Act5C-GAL4* driver and *rpL17A* was used as a loading control. (C) Quantification of the fertility of the male with indicated genotypes (n = 10 individual males/genotype); *p < 0.01 compared with wild type (*WT*). (D) Micrographs of internal male genitalia in *WT* and *Usp14*–mutant flies; protamineB-GFP labeled nuclei of elongated spermatids; arrowheads indicate seminal vesicles. Scale bar = 250 μm.

For more accurate analyses of *Usp14* gene functions, we generated null alleles using CRISPR/Cas9 mediated mutagenesis (Port et al., 2014). Among several indel mutants, we selected the point mutant allele *Usp14*^*Δ32*^ for detailed analysis. In this mutant, insertion of a cytosine after the second codon generates a frameshift and a premature stop codon at 16 base pairs downstream (Fig. 1A). Despite the resulting absence of functional Usp14, *Usp14*^*Δ32*^ null homozygotes were viable but males were sterile, as observed with hypomorph alleles (Fig. 1C, column 4). *Usp14*^*Δ32*^ mutant testes imitated individualization defects in *Usp14*^*f*^ males, indicating no clear differences between phenotypes of null and strong hypomorph alleles. These results suggest that Usp14 is implicated in the coordination of late spermatogenesis.

### *Usp14* mutant male sterility is germline dependent

The male sterile phenotype observed in *Usp14* mutants may exclusively reflect requirements of Usp14 in germline cell lineages and/or testis-forming somatic cells. To identify affected cell types following Usp14 depletion, we used the GAL4/UAS system in Drosophila (Duffy, 2002). Several GAL4 cell lines have specific spatio-temporal expression patterns in fly testis (White-Cooper, 2012). Herein, we used the *bam-Gal4* driver, which had a germline restricted expression pattern in previous studies (Chen and McKearin, 2003). The *Act5c-Gal4* driver was used exclusively to drive expression in somatic cells (White-Cooper, 2012).

*Usp14* specific transgenic RNA interference was induced by the germ line specific *bam-Gal4* driver and resulted in male sterility (Fig. 1C, column 8) with individualization defects that were similar to those in *Usp14* hypomorph and null mutants. Yet, induction of the RNAi with the soma-specific *Act5C-Gal4* driver did not lead to male sterility or phenotypic abnormalities (Fig.1C, column 7). Conversely, the male sterility of *Usp14*^*Δ32*^ homozygotes was rescued by transgenic *Usp14* expression under the control of the *bam-Gal4* driver (Fig.1C, column 10), but not with the somatic *Act5C-Gal4* driver (Fig. 1C, column 9). These results suggest that germline specific expression of *Usp14* is required for proper sperm development.

### *Usp14* mutation disrupts individualization complexes

Germline dependent male sterility and deficient processing of sperm bundles to mature sperms in Usp14 mutant testes suggest that Usp14 facilitates sperm individualization. Cytoskeletal-membrane individualization complexes (IC) mediate sperm individualization in elongated spermatid bundles. IC contains a cluster of 64 actin-rich structures, known as actin cones. During sperm individualization, these move down the bundles and push excess cytoplasm and organelles into so called “cystic bulges” that serve as waste bags.

Concomitantly, spermatids are sheathed in plasma membranes and become individualized and move into seminal vesicles for storage until mating (Noguchi et al, 2006). Severe disturbances of this process lead to individualization defects and stagnation of sperm bundles in the fronts of seminal vesicles.

To determine whether structures of IC are affected in *Usp14* mutants, actin cones were visualized in wild type (WT) and *Usp14*^*Δ32*^ testes after staining with fluorescently labeled phalloidin. In WT testes, triangular shaped actin cones move synchronously (Fig. 2A, upper panels) and actin cones behind the moving IC can only be observed occasionally. However, in *Usp14*^*Δ32*^ testes the majority of ICs lacked synchronous movements of actin cones, and a high proportion of individual actin cones were spread along the sperm bundles (Fig. 2A, lower panels). Hence, Usp14 is likely required for synchronous migration of actin cones, and disruption of IC leads to male sterility. Strikingly similar individualization defects were observed following mutation of the testis-specific proteasome subunit Pros6αT (Zhong and Belote, 2007). Because proteasome interactions of Usp14 are well established in *Drosophila* (Lundgren et al., 2005; Lee et al., 2011), the resemblance of *Pros6αT* and *Usp14* mutant phenotypes suggests that both subunits are required for normal testis-specific proteasome function.

**Fig. 2.**
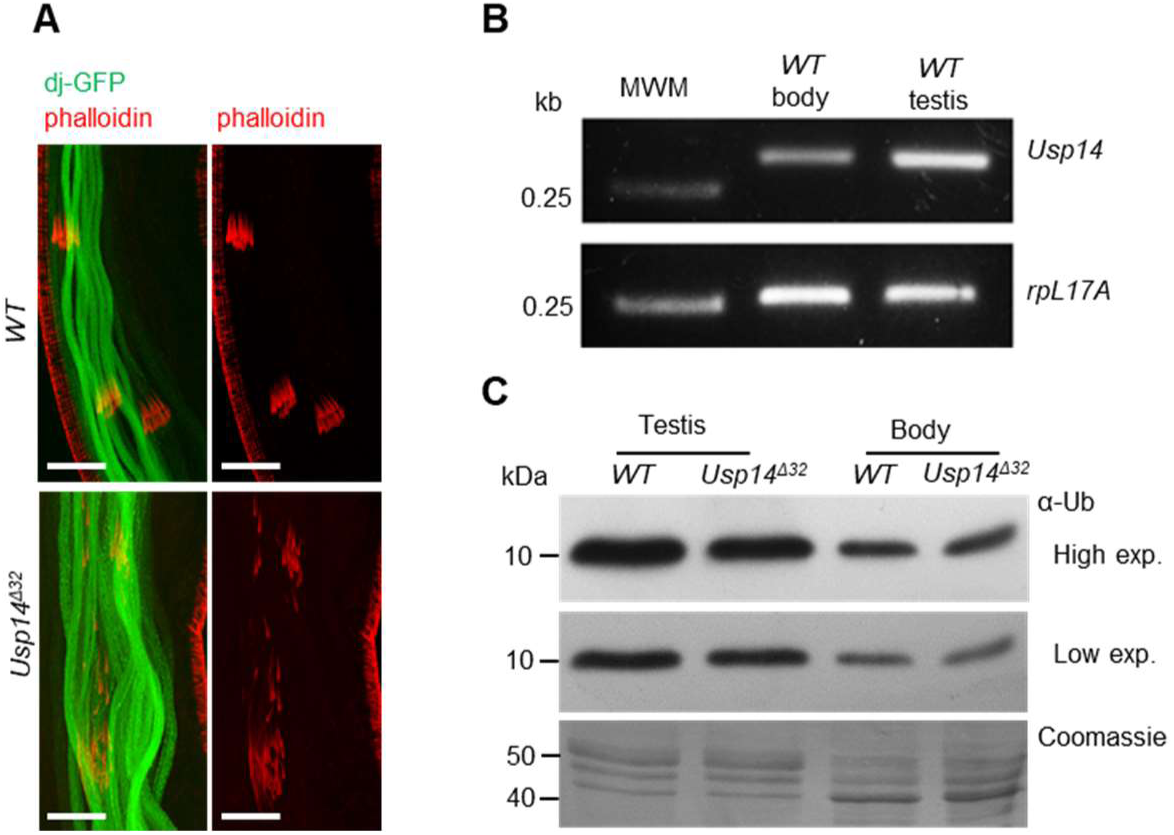
Individualization defects and reductions in monoubiquitin pools in the absence of Usp14; (A) confocal micrographs of sperm bundles; dj-GFP (green) shows sperm tails and phalloidin (red) shows individualization complexes. Scale bar = 5 μm. (B) Expression levels of *Usp14* in testis; Semiquantitative RT-PCR analyses of WT gonadectomized body and testis samples; *rpL17A* was used as a loading control. (C) Monoubiquitin levels in *Usp14* mutant somatic (body) and testis tissues (also shown on Fig. 3B); Western blots show a mono-Ub band at 8.5 kDa. Lanes were loaded with 4 μg aliquots of total protein, as determined using Bradford assays.

### Usp14 is predominantly expressed in testis

Previous studies show that many proteasome subunits and associated proteins have testis-specific orthologues in *Drosophila* (Belote and Zhong, 2009; Ma et al., 2002; Zhong and Belote, 2007), suggesting important roles of the proteasome in *Drosophila* testes. Because Usp14 is a transient but conserved subunit of the regulatory *Drosophila* proteasome particles (Lundgren et al., 2005; Lee et al., 2011), and its mutants show testis-specific phenotypes, we investigated the possibility of differential expression of *Usp14* in testis. To investigate the expression levels of *Usp14*, testis and gonadectomized bodies of WT *Drosophila* were analyzed using semiquantitative RT-PCR experiments. *Usp14* transcripts were more abundant in testis than in other tissues (Fig. 2B). Because accessory glands and other organs were removed from testes before mRNA extraction, somatic cell representation was very low in this sample, including only epithelial cells of the testis surface. With observations of *Usp14* rescue by germline drivers (Fig. 1C), these data indicate that *Usp14* is expressed predominantly in the male germline.

### Loss of Usp14 and reduced free ubiquitin levels in testes

Loss of Usp14 function in yeast and mouse models previously resulted in considerable reductions in monoubiquitin levels (Leggett et al., 2002; Anderson et al., 2005). Usp14 recycles monoubiquitins from protein substrates that are targeted to the proteasome, and prevents their degradation with proteasome targets (Hanna et al., 2003; Anderson et al., 2005). We determined monoubiquitin protein expression levels in WT and *Usp14* mutant testes and gonadectomized body samples using Western blots. We primarily observed significant differences between body and testis monoubiquitin levels. Pixel density analysis of Western blot images demonstrated that the monoubiquitin levels in testis are more than 3-fold higher than in the rest of the body (Fig. 2C, compare lane 1 and 3; and Nagy et al., 2018). The high monoubiquitin concentration in testis suggests that ubiquitination is prevalent during spermatogenesis. Pixel density analyses of immunoblots (Fig. 2C) revealed up to 30% decrease in monoubiquitin concentrations in *Usp14* mutant testes, but no differences in other tissues from WT and *Usp14* mutant flies (see Fig. S1). These data show that monoubiquitin pools in testes are especially sensitive to the loss of Usp14 function, likely contributing to male sterility and individualization defects.

The effects of monoubiquitin deficiencies on spermatogenesis were exacerbated when the *Usp14* mutation was accompanied by a hypomorph allele of the testis-specific ubiquitin gene *Ubi-p63E*. In Drosophila, *Ubi-p63E* is the only polyubiquitin gene that provides newly synthetized ubiquitins in testis (Lu et al., 2013). *Ubi-p63E* null mutant spermatocytes show meiotic arrest with loss of monoubiquitins. The *Ubi-p63E*^*EY*^ hypomorph P element insertion allele, however, has moderate effects on monoubiquitin pools and permits progress of spermatogenesis to the elongated spermatid stage (Lu et al., 2013). After combining *Usp14* ^*Δ32*^ null and *Ubi-63E*^*EY*^ hypomorph mutations, we observed increased severity of the testis phenotype compared with that in single mutants (Fig. 3A). In particular, spermatocytes of the *Usp14*^*Δ32*^*–Ubi-63E*^*EY*^ double mutants had arrested meiosis and no postmeiotic and elongated spermatocytes were observed (Fig. 3A). This phenotype is consistent with the *Ubi-p63E* null phenotype (Lu et al., 2013), with considerable reductions in monoubiquitin levels (Fig. 3B). Perhaps *Usp14* ^*Δ32*^ and *Ubi-63E*^*EY*^ synergistically maintain adequate free monoubiquitin pools in the testis. Loss of *Ubi-p63E* caused severe free ubiquitin shortage leading to early arrest of meiosis, whereas a less severe shortage of ubiquitin in *Usp14 null* mutants manifests in a later stage of spermatogenesis.

**Fig. 3.**
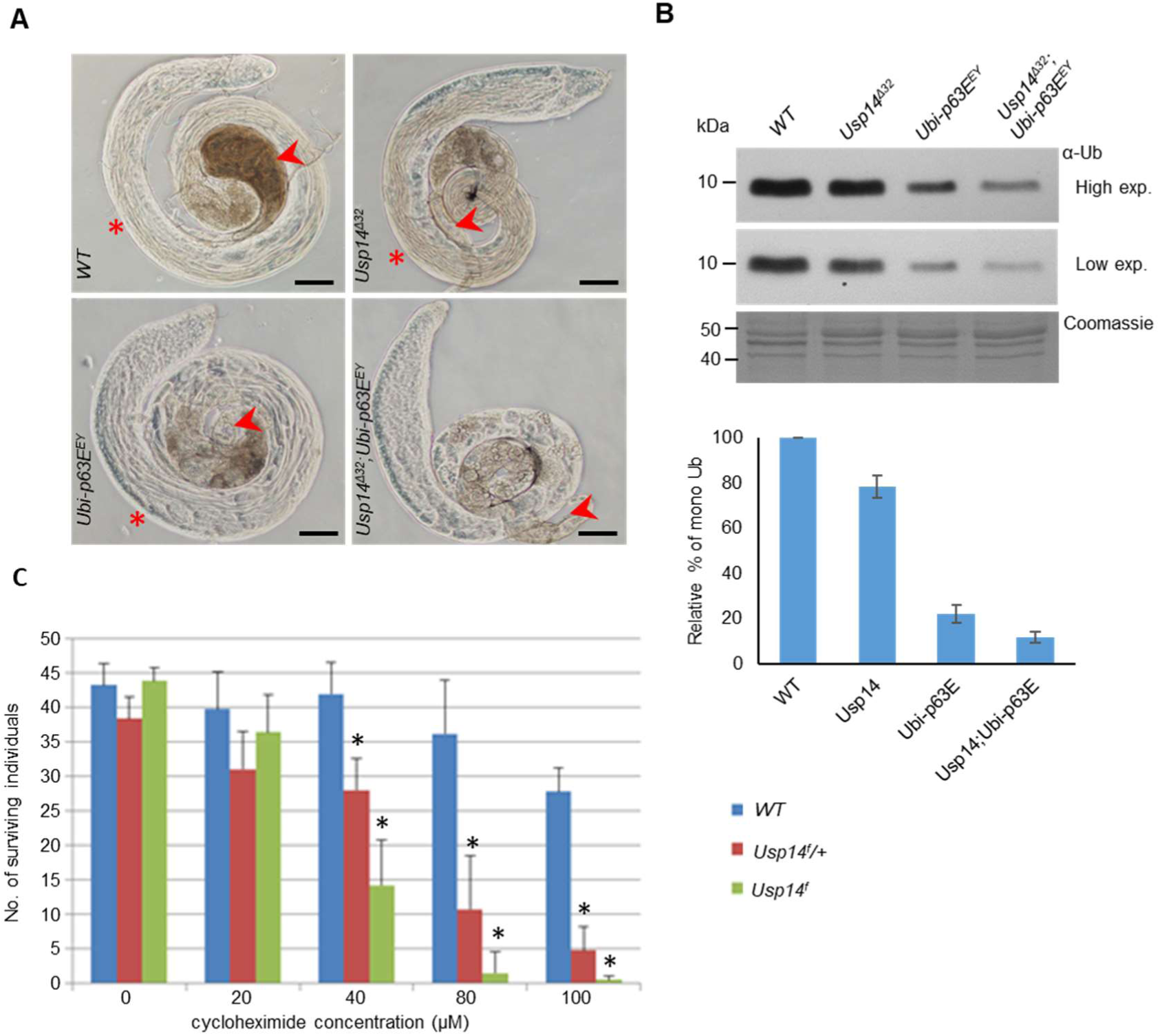
Lack of Usp14 leads to monoubiquitin deficiency. (A) Phase contrast micrographs of testes from males with indicated genotypes; arrowheads show seminal vesicles. Asterisks indicate the presence of elongated spermatids that are absent in Usp14–Ubi-p63E double mutants. Scale bar = 100 μm. (B) Monoubiquitin levels in testis of Usp14 and Ubi-p63E mutants were measured using Western blots with an anti-ubiquitin antibody. After determining protein concentrations using Bradford assays, 4-μg aliquots of total protein were loaded onto gels. The relative percent of the samples (n=4) Monoubiquitin contents in mutants were calculated using data from densitometric analyses and are presented as percentages of that in WT controls; *p < 0.01 compared to WT. (C) Cycloheximide sensitivity of *Usp14* mutants; Data are expressed as means and standard deviations of three independent replicates; *p < 0.01 compared to *WT* data at a given concentration.

### Usp14 mutants are cycloheximide sensitive

The antibiotic cycloheximide is widely used as a eukaryotic protein synthesis inhibitor. It has toxic side effects that are related to rapid depletion of ubiquitin pools in treated cells. Accordingly, expression levels of ubiquitin genes are exceptionally sensitive to this drug (Hanna et al., 2003). Previous studies also identified mutations that reduced intracellular ubiquitin levels and led to cycloheximide sensitivity in yeast and *Drosophila* (Leggett et al., 2002; Chernova et al., 2003; Kovács et al., 2015).

While studying the physiological roles of the Drosophila DUB Usp5, we observed a ubiquitin stress response that had been described in yeast (Hanna et al., 2007). Another study indicated an essential role of Usp14 (Kovács et al., 2015) that was unrelated to its testis-specific function.

Because loss of Usp5 leads to reduced monoubiquitin levels and triggers the expression of Usp14 to counteract ubiquitin degradation and boost ubiquitin recycling, loss of function *Usp14* mutants may be cycloheximide sensitive. Thus to confirm these assumptions, we compared the effects of cycloheximide in WT and hypomorph *Usp14*^*f*^ mutants.

Cycloheximide treatments of *Usp14*^*f*^ homozygote larvae (Fig. 3C) led to cell death in a dose dependent manner, with an LD_50_ of 32 μM. Viability of WT animals was not affected at this concentration. *Usp14f/+* heterozygotes also showed high sensitivity to cycloheximide (LD_50_= 42 μM), indicating that one copy of *Usp14* does not compensate for ubiquitin shortages sufficiently after cycloheximide treatments. These data suggest that Usp14 functions in somatic tissues in situations when the ubiquitin equilibrium is disturbed by reduced ubiquitin expression due to cycloheximide treatment, or by abolished Ub-recycling following elimination of Usp5 (Kovacs et al, 2015). Demand for Usp14 activities in somatic tissues under stressed conditions indicate that Usp14 is not a testis-specific subunit, because it also has roles in somatic tissues. It is more likely that normal spermatogenic processes have high ubiquitin demands and, hence, resemble quasi-ubiquitin stress conditions in which proper functioning of Usp14 and other proteasome subunits is heavily required. Usp14 dependence of spermatogenesis may also be evolutionarily conserved because *Usp14* mutant *ataxia*^*J*^ mice were also sterile and had defects in spermatogenesis (Crimmins et al, 2009). However, loss of Usp14 function also severely affected the neuromuscular system in mice (Wilson et al, 2002; Anderson et al, 2005), suggesting that, unlike in flies, neuronal tissues of more complex organisms are hypersensitive to ubiquitin homeostasis.

Taken together, our data demonstrate pleiotropic roles of *Drosophila* Usp14 deubiquitylase in spermatogenesis and in ubiquitin stress responses. Detailed phenotypic analyses of loss of function alleles show that *Usp14* is essential for normal spermatide differentiation. Accordingly, homozygous *Usp14* mutant males are sterile and have defective sperm individualization processes. In agreement, the phenotype associated with loss of Usp14 function is consistent with its expression pattern in testes.

Finally, individualization defects were strikingly similar to the mutation of the testis-specific proteasome subunit, Pros6αT (Zhong and Belote, 2007). Given that interaction of Usp14 with the proteasome is well established in *Drosophila* (Lundgren et al., 2005; Lee et al., 2011), the resemblance of *Pros6αT* and *Usp14* mutant phenotypes suggest that both subunits are required for normal testis-specific proteasome function.

## MATERIALS AND METHODS

### Drosophila stocks and methods

*Drosophila* lines were maintained on standard yeast/cornmeal medium at 25°C. The PiggyBac element insertion line *Usp14*^*f00779*^ was obtained from Bloomington Drosophila Stock Center (BDSC stock number: 18368). The transgenic RNA interference line *Usp14*^*KK102888*^ was obtained from Vienna Drosophila Resource Center (VDRC stock number: v110227). All genetic markers are described in FlyBase at http://flybase.org. The WT strain *w*^*1118*^ was used for comparisons in all experiments.

To reverse the *Usp14*^*f00779*^ PiggyBac insertion, we crossed stock *en mass* with a PBac transposase source (BDSC:8285) and established two candidate revertant stocks. Restoration of male fertility was observed in both revertants, but only *Usp14* ^*f rv*^ were maintained and used in experiments. CRISPR/Cas9 mediated mutagenesis was used to generate *Usp14*^*Δ32*^ null mutants using previously published tools (Port et al, 2014) with the targeting guide RNA sequence 5’-GCTACCTTTAAATGCGGGCA-3’. Male fertility was tested by placing individual 1-day old males in vials with two 4-day old WT virgin females. After 6 days, adults were discarded and numbers of the eclosed progeny were scored.

### Semiquantitative RT-PCR

Testes of 0–1-day old Drosophila males were dissected in ice cold PBS and tubes containing either 50 testes or 10 gonadectomized males were frozen at −80°C. Total RNA was isolated using Tri Reagent extraction kits (Sigma-Aldrich, USA). RNA samples were treated with RQ1 RNase-Free DNase (Promega Corporation, USA) and were reverse transcribed using Fermentas cDNA synthesis kits. Samples of cDNA were normalized to rpL17A level using PCR with rpL17A forward primer, 5’-GTGATGAACTGTGCCGACAA-3 and rpL17A reverse primer, 5’-CCTTCATTTCGCCCTTGTTG-3’). *Usp14* mRNA expression levels were then determined in 20–25-cycle PCRs with the exon specific *Usp14* RT forward primer 5’-ACGGTGGTGCCCTTCTCC-3’, and *Usp14* RT reverse primer 5’-GGCGCTGTGGTCCTGTTG-3’.

### Fluorescence microscopy

Prior to native observations of GFP fluorescence, testes were dissected and mounted without extensive squashing in PBS and were then directly observed and imaged using an Olympus BX51 upright microscope with phase contrast and UV GFP filters.

Confocal microscopy was performed with testes of 0–1-day old males expressing the sperm tail marker dj-GFP. In these experiments, WT or *Usp14* mutant flies were dissected at room temperature in PBS and were immediately fixed for 20 min in 4% formaldehyde. Testes were stained with AlexaFluor647 conjugated phalloidin (1:400) and DAPI and were then examined under an Olympus FV 1000 confocal microscope.

### SDS-PAGE and Western blot analyses

Testes were dissected in ice cold PBS buffer and were frozen in liquid nitrogen and stored at −80°C until use. Protein samples were then prepared from homogenates of 25 testes in Buffer I F, which contained 100 mM Tris, (pH 7.6), 150 mM NaCl, 1 mM EDTA, 100 mM N-Ethylmaleimide (NEM, Sigma-Aldrich), 20 μM MG132 (Calbiochem) and 1 × EDTA-Free Complete Protease Inhibitor Cocktail (Roche). Homogenates were centrifuged and boiled in 1× Laemli Buffer. Samples containing 4 μg of total protein were loaded onto 14% SDS-acrylamide gels, were electrophoresed and subjected to Western blotting. Protein concentrations were determined using Bradford assays (Bradford, 1976). Monoubiquitin forms of 8.5 kDa were detected using a mouse monoclonal anti-ubiquitin primary antibody (Sigma, U0508, 1:3000 dilution) and a peroxidase-conjugated AffiniPure goat anti-mouse (Jackson Immuno Research, 115-035-003, 1: 30.000 dilution) secondary antibody. Western blots were developed on X-ray films, and were scanned and processed using Image J software.

### Cycloheximide treatment

Cycloheximide (Sigma-Aldrich, USA) was dissolved in EtOH to obtain stock solution. First-instar larvae were collected in vials containing 3.5 ml of standard *Drosophila* medium and were treated with 60 μl aliquots of cycloheximide solution in EtOH. Treatment concentrations were calculated to a final volume of 3.5 ml. Eclosing adults were scored and statistically analyzed using a Microsoft Office Excel software.

## ACKNOWLEDGMENTS

No competing interests were declared by the authors.

This work was funded by grants from the National Research, Development and Innovation Office (OTKA-K116372), Ministry for National Economy of Hungary (GINOP-2.3.2-15-2016-00032). Drosophila stocks obtained from the Bloomington Drosophila Stock Center (NIH P40OD018537) and the Vienna Drosophila Resource Center (VDRC, www.vdrc.at) were used in this study. We also thank Flybase for providing database information (McQuilton et al., 2012).

## REFERENCES

Amerik, A.Y., Hochstrasser, M. (2004). Mechanism and function of deubiquitinating enzymes. Biochim Biophys Acta. 1695, 189–207.

Anderson, C., Crimmins, S., Wilson, J.A., Korbel, G.A., Ploegh, H.L., Wilson, S.M. (2005). Loss of Usp14 results in reduced levels of ubiquitin in ataxia mice. J Neurochem. 95, 724–731.

Belote, J.M., Zhong, L. (2009). Duplicated proteasome subunit genes in Drosophila and their roles in spermatogenesis. Heredity (Edinb). 103, 23–31.

Bradford, M.M. (1976). A rapid and sensitive method for the quantitation of microgram quantities of protein utilizing the principle of protein-dye binding. Anal. Biochem. 72, 248–254.

Chen, D., McKearin, D.M. (2003). A discrete transcriptional silencer in the bam gene determines asymmetric division of the Drosophila germline stem cell. Development. 130, 1159–1170.

Chernova, T.A., Allen, K.D., Wesoloski, L.M., Shanks, J.R., Chernoff, Y.O., Wilkinson, K.D. (2003). Pleiotropic effects of Ubp6 loss on drug sensitivities and yeast prion are due to depletion of the free ubiquitin pool. J Biol Chem. 278, 52102–52115.

Crimmins, S., Sutovsky, M., Chen, P.C., Huffman, A., Wheeler, C., Swing, D.A., Roth, K., Wilson, J., Sutovsky, P., Wilson, S. (2009). Transgenic rescue of ataxia mice reveals a male-specific sterility defect. Dev Biol. 325, 33–42.

Duffy, J.B. (2002). GAL4 system in Drosophila: a fly geneticist’s Swiss army knife. Genesis. 34, 1–15.

Guterman A, Glickman MH. (2004). Deubiquitinating enzymes are IN/(trinsic to proteasome function). Curr Protein Pept Sci. 5, 201–211.

Hanna, J., Leggett, D.S., Finley D. (2003). Ubiquitin depletion as a key mediator of toxicity by translational inhibitors. Mol Cell Biol. 23, 9251–9261.

Hanna, J., Meides, A., Zhang, D.P., Finley, D. (2007). A ubiquitin stress response induces altered proteasome composition. Cell. 129, 747–759.

Kim, J.H., Park, K.C., Chung, S.S., Bang, O., Chung, C.H. (2003). Deubiquitinating enzymes as cellular regulators. J Biochem. 134, 9–18.

Kovács, L., Nagy, O., Pál, M., Udvardy, A., Popescu, O., Deák, P. (2015). Role of the deubiquitylating enzyme DmUsp5 in coupling ubiquitin equilibrium to development and apoptosis in Drosophila melanogaster. PLoS One. 10, e0120875.

Lee, J.H., Park, S., Yun, Y., Choi, W.H., Kang, M.J., Lee, M.J. (2018). Inactivation of USP14 Perturbs Ubiquitin Homeostasis and Delays the Cell Cycle in Mouse Embryonic Fibroblasts and in Fruit Fly Drosophila. Cell Physiol Biochem. 47, 67–82.

Lee, M.J., Lee, B.H., Hanna, J., King, R.W., Finley, D. (2011). Trimming of ubiquitin chains by proteasome-associated deubiquitinating enzymes. Mol Cell Proteomics. 10, R110.003871.

Leggett, D.S., Hanna, J., Borodovsky, A., Crosas, B., Schmidt, M., Baker, R.T., Walz, T., Ploegh, H., Finley, D. (2002). Multiple associated proteins regulate proteasome structure and function. Mol Cell. 10, 495–507.

Lu, C., Kim, J., Fuller, M.T. (2013). The polyubiquitin gene Ubi-p63E is essential for male meiotic cell cycle progression and germ cell differentiation in Drosophila. Development. 140, 3522–3531.

Lundgren, J., Masson, P., Mirzaei, Z., Young, P. (2005). Identification and characterization of a Drosophila proteasome regulatory network. Mol Cell Biol. 25, 4662–4675.

Ma, J., Katz, E., Belote, J.M. (2002). Expression of proteasome subunit isoforms during spermatogenesis in Drosophila melanogaster. Insect Mol Biol. 11, 627–639.

McQuilton, P., St Pierre, S.E., Thurmond, J.; FlyBase Consortium. (2012). FlyBase 101 – the basics of navigating FlyBase. Nucleic Acids Res. 40, D706–14.

Nagy, Á., Kovács, L., Lipinszki, Z., Pál, M. és Deák, P. (2018). Developmental and tissue specific changes of ubiquitin forms in Drosophila melanogaster. PLoS One. 13, e0209080.

Nijman, S.M., Luna-Vargas, M.P., Velds, A., Brummelkamp, T.R., Dirac, A.M., Sixma, T.K., Bernards, R. (2005). A genomic and functional inventory of deubiquitinating enzymes. Cell. 123, 773–786.

Noguchi, T., Lenartowska, M., Miller, K.G. (2006). Myosin VI stabilizes an actin network during Drosophila spermatid individualization. Mol Biol Cell. 17, 2559–2571.

Port, F., Chen, H.M., Lee, T., Bullock, S.L. (2014). Optimized CRISPR/Cas tools for efficient germline and somatic genome engineering in Drosophila. Proc Natl Acad Sci U S A. 111, E2967–76.

Reyes-Turcu, F.E., Ventii, K.H., Wilkinson, K.D. (2009). Regulation and cellular roles of ubiquitin-specific deubiquitinating enzymes. Annu Rev Biochem. 78, 363–397.

Suresh, B., Lee, J., Kim, K.S., Ramakrishna, S. (2016). The Importance of Ubiquitination and Deubiquitination in Cellular Reprogramming. Stem Cells Int. 2016, 6705927.

Tsou, W.L., Sheedlo, M.J., Morrow, M.E., Blount, J.R., McGregor, K.M., Das, C., Todi, S.V. (2012). Systematic analysis of the physiological importance of deubiquitinating enzymes. PLoS One. 7, e43112.

White-Cooper, H. (2012). Tissue, cell type and stage-specific ectopic gene expression and RNAi induction in the Drosophila testis. Spermatogenesis. 2, 11–22.

Wilson, S.M., Bhattacharyya, B., Rachel, R.A., Coppola, V., Tessarollo, L., Householder, D.B., Fletcher, C.F., Miller, R.J., Copeland, N.G., Jenkins, N.A. (2002). Synaptic defects in ataxia mice result from a mutation in Usp14, encoding a ubiquitin-specific protease. Nat Genet. 32, 420–425.

Zhong, L., Belote, J.M. (2007). The testis-specific proteasome subunit Prosalpha6T of D. melanogaster is required for individualization and nuclear maturation during spermatogenesis. Development. 134, 3517–3525.

